# Nonlinear Reconfiguration of Network Edges, Topology and Information Content During an Artifical Learning Task

**DOI:** 10.1101/2020.09.30.321679

**Authors:** James M. Shine, Mike Li, Oluwasanmi Koyejo, Ben Fulcher, Joseph T. Lizier

## Abstract

Network neuroscience has yielded crucial insights into the systems-level organisation of the brain, however the indirect nature of neuroimaging recordings has rendered the discovery of generative mechanisms for a given function inherently challenging. In parallel, neural network machine-learning models have exhibited breakthrough performance in tackling a range of complex problems, however the principles that govern learning-induced modifications to network structure remain poorly understood, in part due to a lack of analytic tools to quantify the dynamics of network structure. While the question of how network reconfiguration supports learning is mirrored in machine learning and network neuroscience, the different contexts of these fields provides a timely opportunity to bring them together synergistically to investigate the problem. Here we combine these two approaches to reveal connections between the brain’s network structure and the emerging network structure of an artificial neural network. Specifically, we train a shallow, feedforward neural network to classify handwritten digits and then used a combination of systems neuroscience and information theoretic tools to perform ‘virtual brain analytics’ on the resultant edge weights and activity patterns of each node. We identify three distinct phases of network reconfiguration across learning, each of which are characterised by unique topological and information-theoretic signatures. Each phase involves aligning the connections of the neural network with patterns of information contained in the input dataset or preceding layers (as relevant). We also observe a process of low-dimensional category separation in the network as a function of learning. Our results offer a systems-level perspective of how artificial neural networks function – in terms of multi-stage reorganization of edge weights and activity patterns to effectively exploit the information content of input data during edge-weight training – while simultaneously enriching our understanding of the methods used by systems neuroscience.

## Introduction

In the human brain, capacities such as cognition, attention, and awareness emerge from the coordinated activity of billions of neurons^1^. Methods that have traditionally been used to map these functions from neuroimaging data were designed to identify ‘activated’, localized regions of the brain that characterize a particular cognitive context^2^. This historical focus on localization has led to a number of key insights about neural function, however it has also made it more challenging to create links between systems-level neural organization and psychological capacities.

A potential means for mapping psychological functions to neural architecture involves the analysis of neuroimaging data from a systems-level perspective^3–5^. By representing the brain as a network of interacting parts, systems neuroscientists are able to characterize high-dimensional datasets in ways that help understand how brain networks process information^6,7^. Across multiple spatial^8^ and temporal^9^ scales, these approaches have revealed a number of systems-level properties of brain organization. A salient example is the measurement of network modularity, which quantifies the extent to which a network is comprised of a relatively weakly inter-connected set of tight-knit sub-modules. Previous whole-brain imaging approaches have shown that network modularity is tightly-linked to performance: modularity increases with learning^5,10^, but decreases during the performance of challenging cognitive tasks^9,11^. These results provide evidence that the network-level topology of the brain is a relevant axis for understanding complex features of human behaviour^12,13^.

Despite their intuitive appeal, current systems-level approaches in human neuroimaging are inherently indirect. For instance, it is currently impossible to map the intact nervous system at the microscale (i.e., cellular) level – instead, we are force to infer structural connectivity indirectly via diffusion weighted imaging^14^, or so-called ‘functional’ connectivity via the similarity of temporal patterns of neural activity or blood flow^15^. Even with access to high resolution images of neural connectivity, we don’t yet have access to generative models that can effectively simulate different patterns of network reconfiguration across contexts. Without these *‘ground truth’* approaches, systems neuroscience is currently stuck at the descriptive level: we can identify consistent changes in network-level reconfiguration as a function of learning^5,10^, or more abstract cognitive capacities, such as working memory manipulation^16^ or dual-task performance^17^, however we have no principled means of translating these observations to interpretable mechanistic hypotheses^18^.

The advent of artificial neural networks (ANNs) in machine learning has the opposite problem: the algorithmic rules for training high-performing networks have been extremely successful, but we don’t yet understand the organizational principles through which reconfigurations of network architectures enable strong performance. Although some of the details of implementation differ^19^, neuroscience and machine learning share some remarkable similarities. For example, the original ANN algorithms were in part inspired by the anatomy of the cerebral cortex^19–21^, and in the case of deep, layered neural networks, both systems share a common property of distributed computation facilitated by complex topological wiring between large numbers of (relatively) simple computational units. Over the last few decades^20^, neural networks have been trained to outperform world experts at complex strategy games, such as Chess and Go^22^. Although the algorithms that are used to train neural network weights are well understood, the manner in which neural networks reconfigure in order to facilitate high levels of classification accuracy remains relatively opaque^2,20,21^. It is this process of adapting a complex network of interacting components to perform a useful task that has escaped a detailed analysis using the established tools of network neuroscience, which themselves have been used to quantify structure–function relationships in the brain for over a decade.

While the question of how network reconfiguration supports learning is mirrored in machine learning and network neuroscience, the different contexts of these fields provides a timely opportunity to bring them together synergistically to investigate the problem^23^. First, we can observe that the process of adapting a complex network of interacting components to perform a useful task is more simply captured in the training of neural networks. Studying this process offers a unique opportunity to study whole-network structure and activity in a controlled setting with a defined learning objective. In this way, we may identify deeper connections between the structure of networks in the brain and in ANNs. For instance, macroscopic human brain networks constructed from multi-region interactions in neuroimaging data demonstrate substantial reconfiguration as a function of task performance: early in the course of learning, the brain is relatively inter-connected and integrated, but this pattern typically gives way to a more refined, segregated architecture as a simple motor skill becomes second-nature^5,10^. Do similar topological changes happen across training iterations of ANNs? Identifying conserved organization properties between a learning brain and a learning ANN could hint at common topological principles underlying distributed information processing.

Furthermore, the synthetic nature of ML networks means that we can directly interrogate the functional signature of specific elements within ML networks as they learn how to classify diverse input examples into a smaller set of outputs classes. While direct access to micro-scale neuronal interconnections is not practically possible using contemporary human neuroimaging approaches, we can directly observe changes in the distributed patterns of connectivity in ANNs over the course of learning. This allows us to investigate how the functional capacities of networks are distributed across their constituent components, which is inherently challenging to study in biological brains. The established tools of network science, as have been applied to quantify structure–function relationships in the brain for over a decade, are perfectly placed for such analysis^24–27^.

Here, we use a network science approach to understand how network reconfiguration supports the performance of ANNs at supervised learning problems. Specifically, we use the tools of systems neuroscience and information theory to analyze a feedforward neural network as it learns to classify a set of binary digits (from the classic MNIST dataset^28^).

The fact that this classic dataset is so well understood enables us to more clearly interpret how network reconfiguration supports learning. Importantly, given the similar principles at play in more complex neural networks, which either alter architectural^29^ or nodal features^30^ while keeping basic principles of training intact, we anticipate that any conclusions gleaned from the study of extremely simple network architectures can be used as the basis of future interrogation of more complex architectures. If we find similarities between network properties in two classic and high-performing distributed information processing systems — the brain and ANNs — it could provide hints as to more general principles of the properties of underlying network architectures that facilitate efficient distributed information processing.

In particular, we were interested in whether the topology of the neural network over the course of learning mirrored patterns observed in the analysis of fMRI networks in human participants^13^. By tracking functional networks derived from fMRI data over the course of 10 sessions in which participants learned to map visual stimuli to motor responses, it was observed that effective learning was associated with an increase in network modularity^13^, *Q*, which quantifies the extent with which the network can be clustered into tight-knit communities with relatively sparse connections between them^13^. A plausible explanation of these findings is that structural connections within the modularized regions increased their strength over the course of learning, however technological limitations make this inference challenging. Fortunately, we can leverage the full observability and tractability of feedforward neural networks to directly test these ideas *in silico*, and indeed contribute to the cause of “explainable AI”^19–21^.

By tracking the topology of the MNIST-trained network over the course of training, we partially confirmed the original hypothesis of increasing segregation as a function of learning, however our analysis identified a more subtle temporal partition. Early in learning, training reconfigured the edges of the network so that they are strongly aligned with information-rich regions of the nodes in up-stream layers of the network, but in a manner that did not alter the global topology of the network (i.e., the edges did not become more modular during the first phase). Following this initially topologically silent phase, the network then entered a second phase characterized by a rapid increase in modularity that was coincident with large gains in classification accuracy. Later in learning, network-activity patterns reconfigured to a slightly less modular state that maximized digit category separation in a low-dimensional state space estimated from the activity patterns of the nodes within the network. Our results provide foundational understanding of how ANN network activity and connectivity evolves over the course of learning that simultaneously informs our understanding of both systems neuroscience and machine learning.

## Results

### Feed-Forward Neural Network Construction and Training

We applied systems neuroscience and information theoretic methods to analyze the structure of a feed-forward neural network as it was trained to rapidly classify a set of ten hand-written digits (Modified National Institute of Standards and Technology [MNIST] dataset^28^). The ANN was trained across 100,000 epochs with stochastic gradient descent, however we only present a subset of epochs in order to demonstrate the key patterns observed in the dataset – specifically, we analyze a total of 64 epochs: the first 30; every 10 epochs to 100; every 100 epochs to 1000; every 1000 epochs to 10,000; and every 10,000 epochs to 100,000. Although a neural network with a single hidden layer is theoretically sufficient for high performance on MNIST^28^, neural networks with more hidden layers provide benefits of both computational and parameter efficiency^31^. For the sake of simplicity, we chose a relatively basic network in which edge weights and nodal activity patterns could be directly related to performance.

With these constraints in mind, we constructed a feed-forward network with two hidden layers – a 100-node hidden layer (HL1) that received the 28 × 28 input (pixel intensities from the MNIST dataset) and a 100-node hidden layer (HL2) that received input from HL1 – and a 10-node output layer (Fig. 1A). The edges between these layers were given unique labels: edges connecting the input nodes to the first hidden layer were labelled as α edges (dark blue in Fig. 1A); the edges connecting the two hidden layers were labeled as β edges (orange in Fig. 1A); and the edges connecting the second hidden layer to the readout layer were labelled as γ edges (dark green in Fig. 1A). A clear difference between the topology of the ANN and standard approaches to analysing neuroimaging data is that the mean of the absolute value of edge weights from all three groups increased nonlinearly over the course of training in the ANN, whereas typical neuroimaging analyses normalize the strength of weights across cohorts.

**Figure 1.**
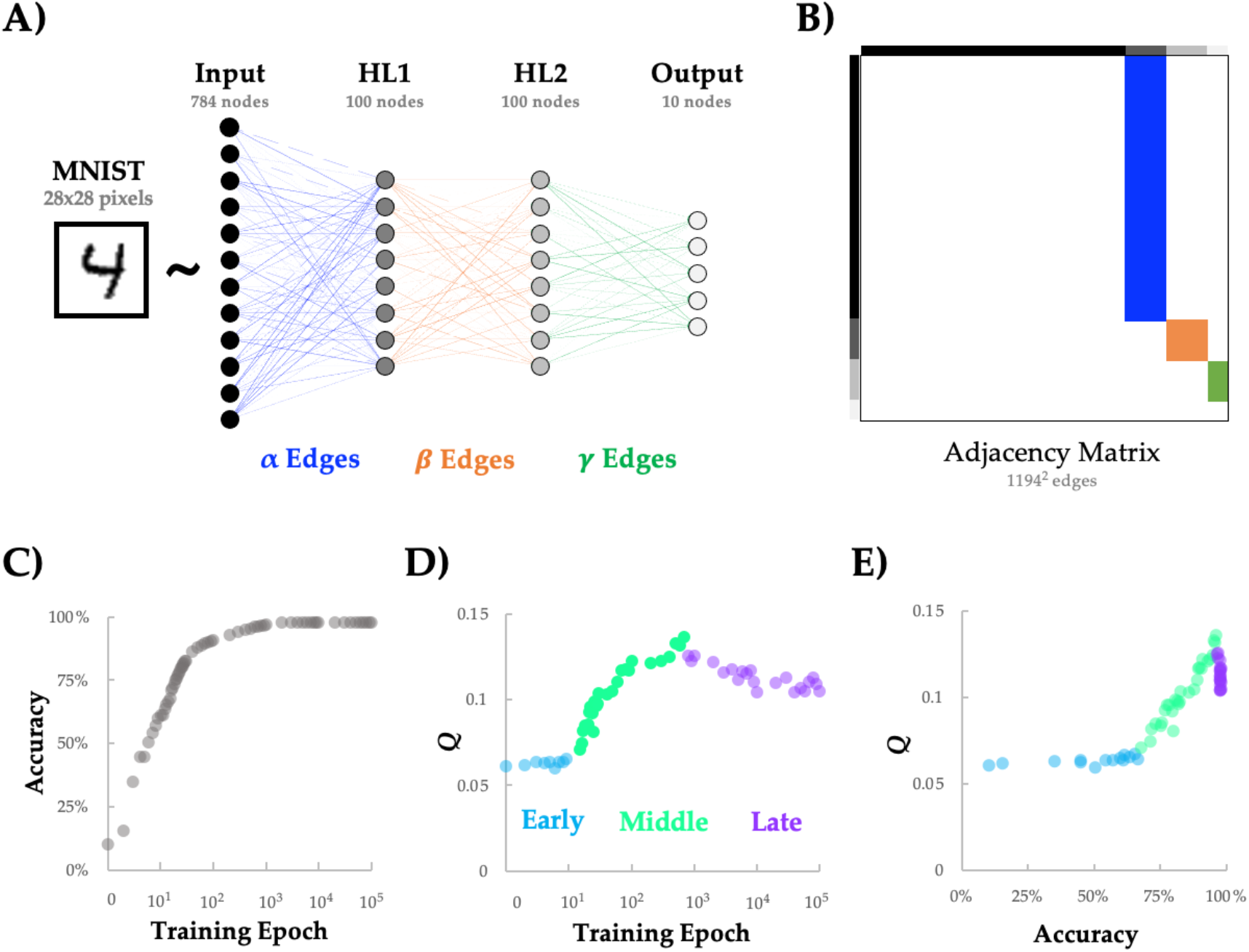
A feed-forward neural network exhibits three topologically distinct periods of reconfiguration throughout learning the MNIST dataset. A) A large (60,000 item) corpus of hand-drawn digits from the MNIST database (28 × 28 pixel array with 256 intensity values per pixel) were vectorized and entered into a generic feed-forward neural network with two hidden layers – a 100-node layer (HL1) that received the 28 × 28 input and a 100-node layer (HL2) that received the input from HL1 – and a 10-node output layer (argmax); B) the edges connecting the input ➔ HL1 (dark blue; α), HL1 ➔ HL2 (orange; *β*) and HL2 ➔ output (dark green; *γ*) were embedded within an asymmetric weighted and signed connectivity matrix; C) Classification accuracy increased rapidly in the early stages of training, with an asymptote after ~100 training epochs; D) Network modularity (Q) was naturally grouped into three separate periods: an Early period (light blue; epochs 1-14) that was relatively static, a Middle period (light green; epochs 15-700) with a rapid increase in Q, and a Late period (light purple; epochs 800-10,000) in which *Q* diminished, albeit not to initial levels. E) classification accuracy showed a non-linear relationship with *Q*: initial increases in accuracy were independent of *Q* (light blue), after which there was a positive linear relationship between accuracy and *Q* (Pearson’s *r* = 0.981; light green), and finally a sustained drop in *Q*, as accuracy saturates in the later periods of learning (light purple). For clarity, only a subset of the 100,000 epochs are presented here.

### The Topological Properties of a Feed-Forward Neural Network During Training

It has previously been suggested that the concept of modularity (i.e., *‘Q’*) may be employed to improve the design of deep neural-network architecture in various ways^32,33^. Non-trivial modular structure is a pervasive feature of complex systems^25,27^, and has been shown to increase as a function of learning in neuroimaging experiments^5,10^. Based on this similarity, we hypothesized that Q should increase as a function of training on the MNIST dataset and should reflect improvements in classification accuracy. To test this prediction, we required a means for translating the edges of the neural network into a format that was amenable to network science approaches (i.e., a weighted and directed adjacency matrix). To achieve this, we created a sparse node × node matrix, and then mapped the α (Input–HL1), *β* (HL1–HL2) and *γ* (HL2–output) edges accordingly, yielding the adjacency matrix shown in Fig. 1B.

With the network edge weights framed as a graph, we applied methods from network science to analyse how its complex topological structure changed as the ANN was trained to classify the MNIST dataset (Fig. 1C). We applied used the Louvain algorithm to estimate *Q* from the neural network graph at each training epoch. Variations in network modularity across training epochs are plotted in Fig. 1D and reveal three distinct periods of: approximately constant *Q* (‘Early’; training epoch 1-9; data points 1-9 in Fig. 1D), followed by increasing *Q* (‘Middle’; training epoch 10-8,000; data points 10-55 in Fig. 1D), and finally decreasing *Q* (‘Late’; training epoch 9,000-100,000; data points 56-64 in Fig. 1D). Early in training, there was a substantial improvement in accuracy without a noticeable change in *Q* (light blue in Fig. 1E). In the Middle period, we observed an abrupt increase in *Q* (light green in Fig. 1E) that tracked linearly with performance accuracy (*r* = 0.981, *p_PERM_* < 10^−4^, permutation test). Finally, in the Late training period, *Q* began to drop (Fig. 1E; light purple). These results demonstrate that the modularity of the neural network varies over the course of training in a way that corresponds to three different types of behavior with respect to the network’s classification performance.

### Early Edge-weight Alteration is Concentrated on Informative Inputs

The fact that *Q* didn’t change early in training, despite substantial improvements in accuracy, was somewhat surprising. This result was made even more compelling by the fact that we observed substantial edge-weight alteration during the Early period, however with no alteration in modularity. To better understand this effect, we first created a difference score representing the absolute value of edge changes across each pair of epochs in the Early phase (i.e., the blue epochs in Fig. 1D/E). We then calculated the grand mean of this value across the first epoch (i.e., one value for each of the 784 input dimensions, summed across all α edge weights associated with each input node in the Input layer), and then reshaped this vector such that it matched the dimensions of the Input data (i.e., 28^2^ pixels). We found that the α edge weights that varied the most over this period were located along the main stroke lines in the middle of the image (e.g., the outside circle and a diagonal line; Fig. 2A).

**Figure 2.**
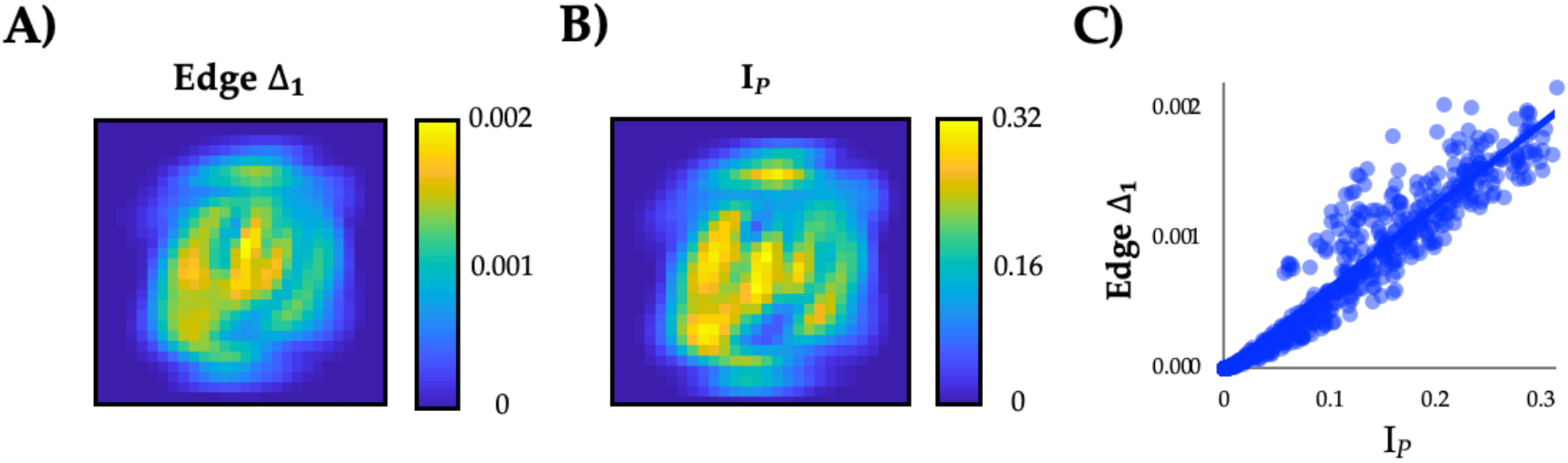
Topologically silent alterations in network edges during the Early period of training. A) although network modularity was static in the Early period, the standard deviation of changes in edge strength, Edge Δ_1_, in the first hidden layer of the network did change substantially over the course of the Early training period (first 10 epochs; cf. Fig. 1C); B) Pixel information, I*_P_* = MI(pixel,class); C) We observed a strong positive correlation between Edge Δ_1_ and I_*P*: *r*_ = 0.965 (p < 1^−10^).

Similar to the manner in which an eye saccades to a salient target^34^, we hypothesized that the feed-forward network was reconfiguring early in training so as to align with the most informative regions of the input space. To test this hypothesis, we binarized the pixel activity across the 60,000 items from the training set, with a threshold that varied across each pixel so as to maximize the mutual information (MI) that the binarized pixel provides about the class (i.e., the digit), and then calculated the information held by each pixel (I*_P_*: MI(pixel,class); Fig. 2B). We observed a clear, linear correspondence between I*_P_* and the edges that reconfigured the most during the Early period (Fig. 2C; *r* = 0.965, *p_PERM_* < 0.0001). The effect remained significant for edge changes in the Middle (r = 0.874) and Late (r = 0.855) periods, however the effect was significantly strongest for the Early period (Z = 16.03, p < 0.001)^35^. This result indicates that the network was adjusting to concentrate sensitivity to class-discriminative areas of input space, which we demonstrate occurs via the reconfiguration of edge weights relating to the most class-discriminative areas of the input space.

### Topological Segregation During the Middle Period of Learning

Following the initial period of learning, we observed a substantial increase in network modularity, *Q*, that rose linearly with improvements in classification accuracy (Fig. 1C, green). To better understand how node-level network elements reconfigured during the Middle period, we computed two metrics for each node that quantify how its connections are distributed across network modules: (i) module-degree *z*-score (MZ); and (ii) participation coefficient (PC)^36^. MZ and PC have together been used characterize the cartographic profile of complex networks: MZ measures within-module connectivity, and PC measures between-module connectivity and thus captures the amount of inter-regional integration within the network (see Methods for details; Fig. 3A)^36^. These statistics have been used previously in combination with whole-brain human fMRI data to demonstrate a relationship between heightened network integration and cognitive function^11,37^, however the role of integrative topological organization is less well understood in ANNs. Importantly, the calculation of both MZ and PC relies on the community assignment estimated from the Louvain algorithm, and hence affords a sensitivity to changes in network topology over the course of training.

**Figure 3.**
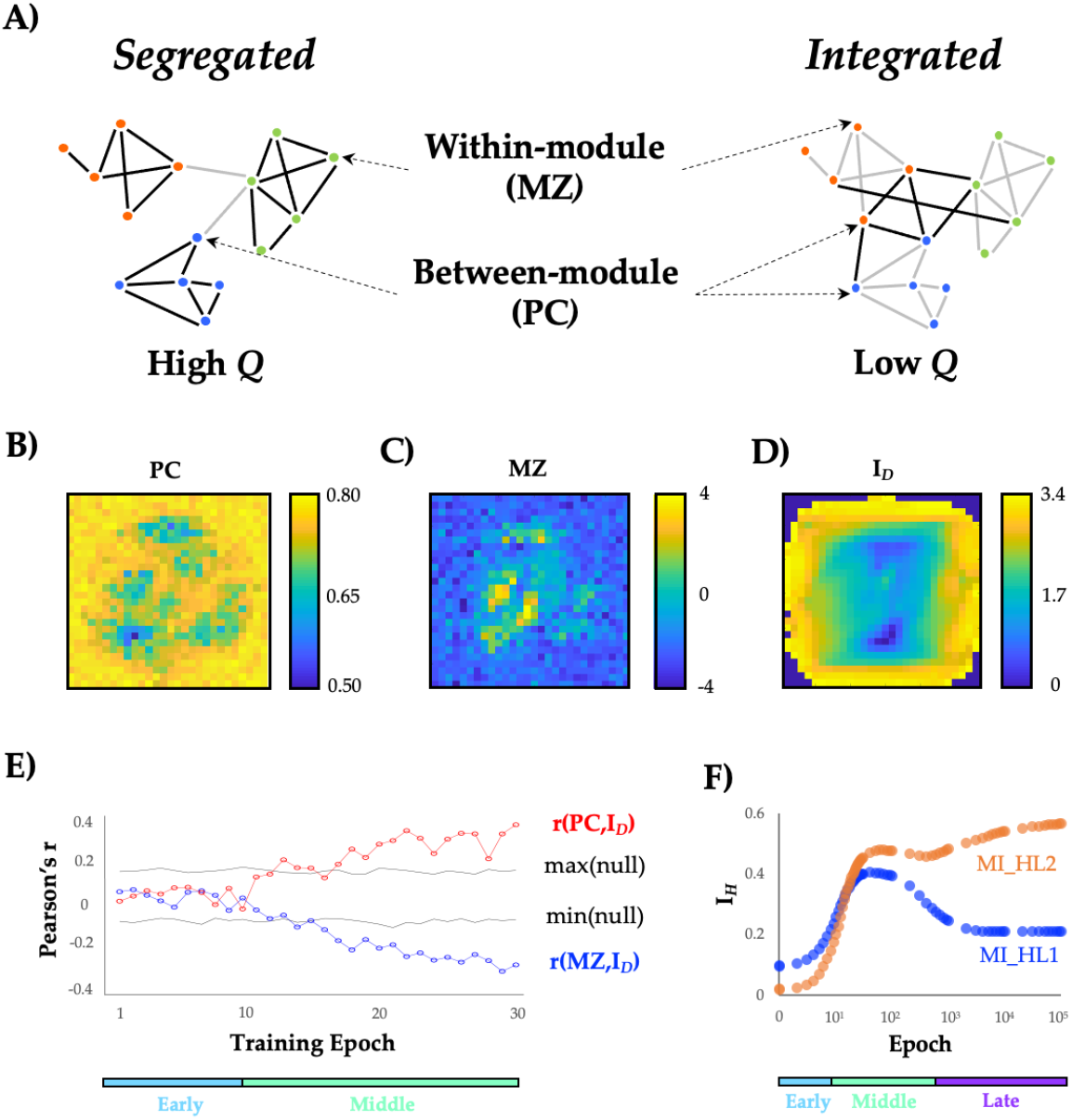
Topological changes during the Middle period. A) a schematic depiction of two topological extremes: on the left is a segregated network, with tight-knit communities that are weakly-interconnected – this network would be characterized by high *Q*, and would have more nodes with high module degree *z*-score (MZ) than nodes with high Participation Coefficient (PC); on the right is an integrated network, which has stronger connections between nodes in different communities, and hence a lower *Q*, and more nodes with high PC than nodes with high MZ; B) participation coefficient (PC) of Input layer nodes at training epoch 30; C) module degree *z*-score (MZ) of Input layer at training epoch 30; D) Digit information, I*_D_* = MI(pixel*On*,class); E) Pearson’s correlation, *r*, between I*_D_* and PC (red) and MZ (blue) across first 30 training epochs. Black lines represent the upper and lower bounds (95th and 5th percentiles) of a permuted null dataset (10,000 iterations) and coloured bars represent learning periods; F) I*_H_* = MI(node,class) for HL1 (blue) and HL2 (orange) edges – note that both subnetworks increase I*_H_* during the Middle period, but that the Late period dissociates the two subnetworks.

Using this cartographic approach^36^, we were able to translate the edge weights in the network into values of PC and MZ for each node of the network for each epoch of training. Figures 3B and 3C show the PC and MZ values for the nodes in the Input layer (i.e., the topological properties of the α edges) at training epoch 30, which was indicative of the patterns in the Middle period. PC was associated with a relatively ‘patchy’ appearance around the main stroke areas, suggestive of a distributed topological coverage of the input space, as well as high values on the edges of the input space (Fig. 3B). In contrast, MZ values were far more centrally concentrated, indicative of local hubs within network communities around the main stroke areas of the inputs space (Fig. 3C). Overall, PC and MZ mapped onto different locations in the input space, and hence were somewhat negatively correlated when data were pooled across all epochs (*r* = –0.107; *p* = 3 × 10^−89^). We hypothesized that these changes in MZ and PC were indicative of a topological reconfiguration of the input layer of the ANN to align network hubs with key aspects of the input stream, being the main stroke areas here.

To test this hypothesis, we related the PC and MZ for each node of the network across all epochs of training to a statistic, I*_D_*: MI(pixelOn,class), which computes the amount of information available in each pixel of the input space when that pixel is active (Fig. 3D). In contrast to the average information I*_P_* held by the pixel about the class, I*_D_* is a partial information, quantifying how informative each pixel is for tracking multiple different digit classes *only* when the pixel is active (pixelOn). High values of I*_D_* imply that the recruitment of the pixel is associated with a reduction in uncertainty (i.e., an increase in information) about the digit. As detailed in the Methods, I*_P_* (Fig. 2B) is negatively correlated to I*_D_* (Fig. 3D) and dominated by samples when the pixel is inactive.

We observed a significant positive correlation between I*_D_* and MZ that emerged towards the end of the Middle period (Fig. 3E). Specifically, we observed a dissociation in the input layer (Fig. 3E) during the Middle period, wherein I*_D_* was positively correlated with PC (max *r* = 0.396, *p_PERM_* < 10^−4^), but negatively correlated with the module degree *z*-score (max *r* = −0.352, *p_PERM_* < 10^−4^). In other words, the topology of the neural network reconfigured so as to align highly informative active pixels with topologically integrated hubs (nodes with higher PC). Whilst these pixels are less commonly active, they are highly informative of class when they are active (high *I_D_*), suggesting that the pixel being active requires the network to send information about such events to many downstream modules. By contrast, more segregated hubs (nodes with higher MZ) are more likely to be associated with higher I*_P_*, being nodes that are more informative on average of digit class (and tending to be more highly informative when inactive). This may indicate that the network is reconfiguring so as to organize sets of informative nodes into modules in a way that supports the creation of higher-order ‘features’ in the next layer. In neuroscience, nodes within the same module are typically presumed to process similar patterns of information^13^, suggesting that the topology of the neural network studied here may be adjusting to better detect the presence or absence of low-dimensional features within the input space.

### Inter-layer Correspondence

Given that the same gradient descent algorithm used to train the network was applied consistently across all layers of the network, we predicted that the same principles identified in the input layer should propagate through the network, albeit to the abstracted ‘features’ captures by each previous layer. Similar to the manner in which a locksmith sequentially opens a bank vault, we hypothesized that each layer of the neural network should align with the most informative dimensions of its input in turn, such that the information could only be extracted from an insulated layer once a more superficial layer was appropriately aligned with the most informative aspects of its input stream. To test this hypothesis, we investigated how the mutual information I*_H_*: MI(node,class) between each node’s activity and the digit class evolved across training epochs. (Note that I*_H_* is equivalent to I*_P_* but computed for hidden layer nodes rather than inputs). As shown in Fig. 2F, mean MI within both hidden layers 1 (MI*_HL1_*) and 2 (MI*_HL2_*) increased during the first two epochs, but then diverged at the point in learning coinciding with the global decrease in modularity, *Q* (cf. Fig. 1D). Crucially, despite the decrease in MI*_HL1_* there was still an increase in MI*_HL2_*, suggesting that the Layer 2 nodes are improving their ability to combine information available in separate individual Layer 1 nodes to become more informative about the class. This suggests that Layer 1 nodes specialize (and therefore hold less information overall, lower MI*_HL1_*) in order to support the integration of information in deeper layers of the neural network (increased MI*_HL2_*).

### Validation with the eMNIST Dataset

In summary, in studying the topological reconfiguration of an ANN during training on the MNIST dataset, we observed three distinctive periods of adjustment, which play different roles in augmenting the distributed information processing across the network to capture class-relevant information in the input data. To better understand the generalizability of these findings, we trained a new feed-forward neural network (identical in architecture to the original network) on the eMNIST dataset^52^. The eMNIST dataset is similar to MNIST, but uses hand-written letters, as opposed to numbers. Although learning was more protracted in the eMNIST dataset (likely due to the increased complexity of the alphabet, relative to the set of digits), we observed similar changes in network structure across training as reported above for the MNIST dataset. Specifically: (i) the network shifted from integration to segregation; (ii) layers reconfigured in serial; and (iii) nodal roles (with respect to inferred network modules) were similarly related to class-relevant information in individual pixels (Fig. S1). These results suggest that the insights obtained from the MNIST analysis may represent general topological features of efficient distributed information processing in complex systems.

### The Late Period is Associated with Low-Dimensional Pattern Separation

Next, we investigated whether the extent to which the nodal topology of the networks trained on the two datasets differed (i.e., whether different regions of the input space had higher or lower PC and MZ) was proportional to the most informative locations of the input space in each dataset (ΔI*_D_*). Specifically, the difference in the pattern of an input node’s edges across inferred network modules between the eMNIST and MNIST datasets (ΔPC) was correlated with the difference in image input characteristics between the two datasets (ΔI*_D_* vs. ΔPC: *r* = 0.301, *p_PERM_* < 0.0001; ΔI*_D_* vs. ΔMZ: *r* = −0.247, *p_PERM_* < 0.0001). This result provides further confirmation that neural networks learn by reorganizing their nodal topology into a set of periods that act to align network edges and activity patterns with the most informative pixels within the training set.

We found that pixels demonstrated unique roles across learning with respect to the emerging modular architecture of the neural network, and that these roles were shaped by their class-relevant information. As edge weights were reconfigured across training, we observed that standard deviation of changes in outgoing edge strength from a node (i.e., Edge Δ1) increases for highly informative inputs (i.e., high I*_P_*; Fig. S1D for eMNIST corresponding to Fig. 2C for MNIST). As these weights change, they alter the activity of each of the nodes in the hidden layers, which ultimately pool their activity via modules to affect the class predictions, which are read out based on the activity of the final output layer. So how do the changes in edge weight translate into nodal activity? Based on recent empirical electrophysiological^38^ and fMRI^39^ studies, we hypothesized that the activity patterns would be distributed across the neural network in a low-dimensional fashion. Specifically, by way of analogy to the notion of manifold untangling in the ventral visual system^40^, we predicted that across training, the high-dimensional initial state of the system (i.e., random edge weights) would become more low-dimensional as pixel-pixel redundancies were discovered through the learning process.

To test this hypothesis, we used dimensionality-reduction^41^ to analyze the ‘activity’ of all of the nodes within the neural network, across the different training epochs. Here, activity was defined as the sum of weighted inputs from inputs or earlier layers of the network, after having been filtered through an activation function. We applied PCA^42^ to the nodal activity across all four layers of the feed-forward network – i.e., the Input, HL1, HL2 and Output nodes – which were first standardized and then either concatenated (to calculate the dimensionality of the entire process) or analyzed on an epoch-to-epoch basis (to calculate the effect of training; see Methods for details). The concatenated state-space embedding was relatively low-dimensional (120/994 components, or 12.2%, explained ~80% of the variance) and the pixel-wise loading of each of the top eigenvalues (λs) for the Input layer (Fig. 4A) was correlated with both *I_P_* and *I_D_* statistics used in the prior analyses (I*_P_* – λ_1_: *r* = 0.218, *p* < 10^−4^; λ_2_: *r* = 0.189, *p* < 10^−4^; λ_3_ = 0.158, *p* < 0.0001; and I*_D_* – λ_1_: *r* = 0.338, *p* < 10^−4^; λ_2_: *r* = 0.123, *p* < 10^−4^; λ_3_: *r* = 0.062, *p* = 0.08), suggesting a direct correspondence between class-relevant information in the input space and the low-dimensional embedding. Crucially, test trials that were incorrectly classified (at Epoch 10,000, though results were consistent for other epochs) were associated with lower absolute loadings on the ten most explanatory EVs (EV_1-10_; Figure S3; FDR *p* < 0.05). These results are tangentially related to recent empirical neuroscientific studies that employed dimensionality reduction on electrophysiological^38^ and fMRI data^39^ to show that learning and cognitive task performance are typically more effective when constrained to a low-dimensional embedding space.

**Figure 4.**
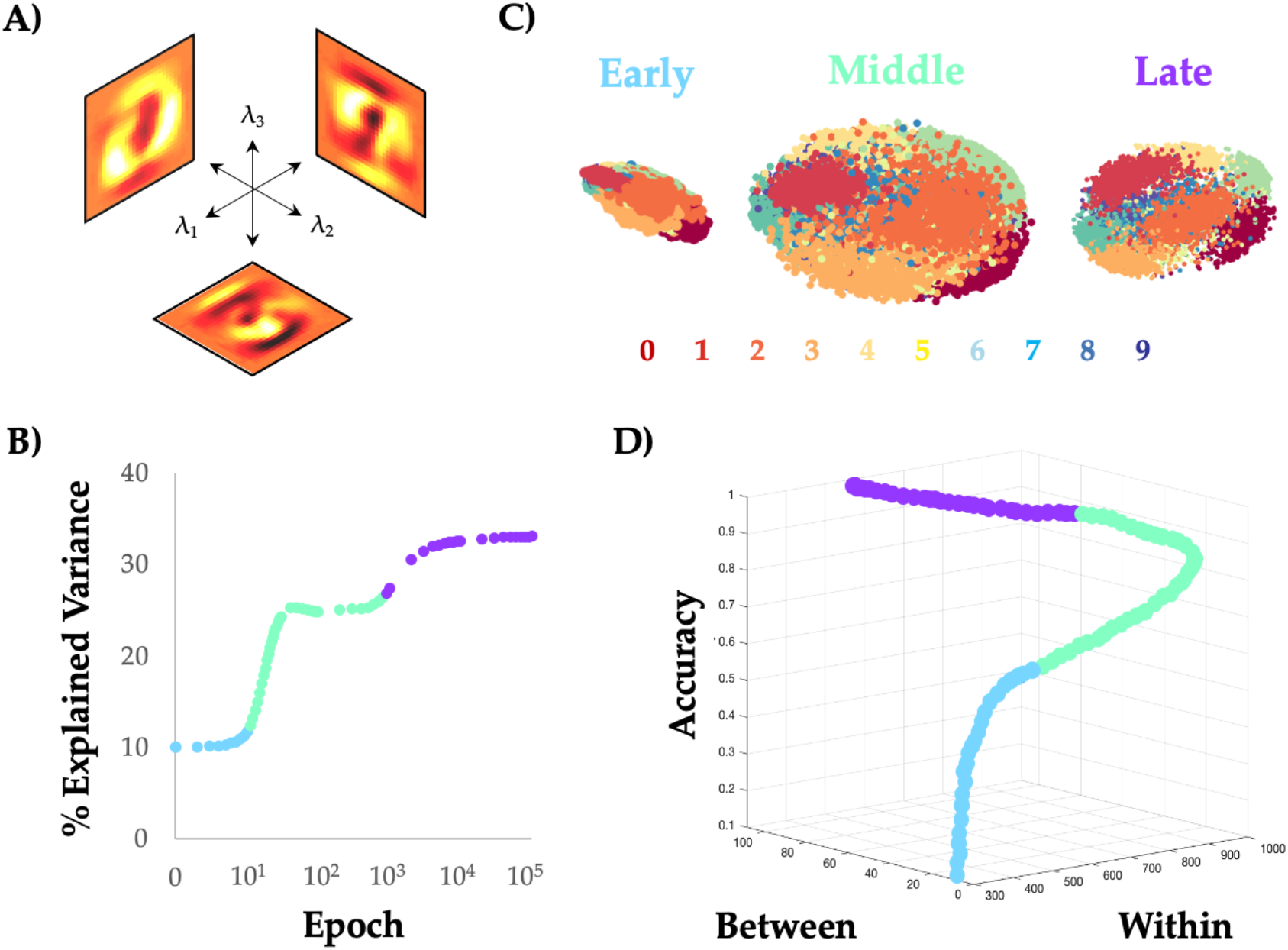
Unravelling the manifold: low-dimensional projections of feed-forward neural network activity during MNIST training reveal category-specific untangling. A) The first three principal components (eigenvalues 1-3: λ_1_/ λ_2_/ λ_3_) of the Input nodes; B) The percentage of variance explained by EV_1_, when the PCA was fit on data from each training epoch separately; C) 3D scatter plot of the items from the training set during three different periods: during the Early period (Epochs 1–10), the topological embedding of the different digits showed substantial overlap, which is reflected in the low between-category distance (i.e., distance between mean of each digit); in the Middle period (Epochs 11–300), the embedding showed a relative expansion in the low-dimensional space; and during the Late period (Epochs 300+), the distance within each category dropped dramatically; D) 3D scatter plot of between-category and within-category distance, along with training accuracy – note that maximal accuracy is associated with increases in both within- and between-category distance.

By conducting a PCA on each epoch in turn, we found that training was associated with a nonlinear alteration in the amount of variance explained by the top 10 PCs (Vd^10^), and that these changes aligned well with the topologically identified periods (Fig. 4B and Movie S1). The network began in a relatively high-dimensional configuration, consistent with the random initiation of nodal activity. During the Early period (light blue in Fig. 4B), as the edge weights reconfigured to align with I*_P_* (Fig. 3D), Vd^10^ remained relatively high. During the Middle period (light green in Fig. 3B), there was a sharp reduction in Vd^10^, however the dimensionality collapse was diminished mid-way through the period. The Late period (purple in Fig. 3B) was associated with a mild reduction in Vd^10^. Interestingly, heightened levels of training led to a tighter correspondence between nodal topological signatures (PC/MZ calculated separately at each epoch) and the principal component loadings of nodes in the Input layer (Fig. S2), suggesting that the topology of the neural network reconfigured over training to better map onto a low-dimensional sub-space that concentrated class-relevant information in the training dataset.

Organising an information processing system within the constraints of a relatively low-dimensional architecture (i.e., dimensions << nodes) can confer important computational benefits^41^. For instance, previous theoretical work in systems neuroscience has argued that the ventral visual stream of the cerebral cortex is organised so as to ‘untangle’ different inputs streams into highly informative categories^40^. Here, ‘untangling’ refers to the ability of the system to effectively separate inputs along different categorical dimensions (e.g., distinguish a well-known face from that of a stranger), while still retaining sufficient information in the signal such that higher-order classifications of the same data are still possible (e.g., recognising a well-known face in a unique orientation). Interestingly, the same concept has been used to explain the function of both the visual system^40^ and effective decision making^43^, and may underpin the functionality of convolutional neural networks trained on naturalistic images^24^. In the context of our PCA analysis, ‘untangling’ could be envisaged as alterations in the way that the activity of the network reflecting different digit categories is embedded within the network’s state space: the loadings onto different categories in the untrained network should be relatively overlapping (i.e., ‘tangled’), but should become less overlapping (i.e., ‘untangled’) as the network learns to effectively categorise the inputs into distinct digits.

Analysing our data from this vantage point, we found that the increase in topologically rich, low-dimensionality was associated with a relative ‘untangling’ of the low-dimensional manifold (Fig. 4C): the Middle period was associated with a general expansion in the low-dimensional embedding distance Within categories (light green in Fig. 4D), which then allowed the system to both expand Between categories and contract within Categories during the Late period of learning (purple in Fig. 4D). This ultimately had the effect of boosting classification accuracy. Indeed, the contraction of the within category embedding distance – which takes place first – co-occurs with the drop of MI*_HL1_*, with the following expansion of Between category distance co-occurring with the increase in MI*_HL2_*. At the sub-network level, the activity on nodes in HL2 was substantially more low-dimensional than HL1 (Fig. S4), further expanding on the notion that different computational constraints are imposed on neural networks, depending on the depth of network layers. Overall, these results confirm the presence of manifold ‘untangling’ in a simple, feed-forward ANN, and hence provide a further link between the way that both synthetic and biological neural networks learn how to classify visual inputs.

## Discussion

In this work, we used information theoretic and network science tools to study the topological features of a training neural network that underly its performance on supervised learning problems. We found many similarities between the topological properties of the brain and ANNs – two systems known for efficient distributed information processing. In the training ANN, we observed three distinctive periods of topological reconfiguration, in which changes in edge-strength (Fig. 2), topology (Fig. 3) and low-dimensional activity (Fig. 4) showed a striking correspondence with class-relevant information in the input data. The results of our study help to both validate and refine the study of network topology in systems neuroscience, while also improving our understanding of how neural networks alter their structure so as to better align network edges and the topological signature of the network with the available streams of information delivered to the input nodes.

The reconfiguration of network edges occurred in three distinct periods over the course of learning. In the first (relatively short) period (light blue in Fig. 1C), the weights of edges in the first layer were adjusted in proportion to the amount of class-relevant information related to each node (i.e., I*_P_*). This process achieved “easy” increases in accuracy, with very little alteration to the modular structure of the network, and only modest increases in information held by the hidden layer nodes about the digit class. The next, somewhat more protracted period (light green in Fig. 1C) involved a substantial topological reconfiguration that resulted in large increases of network modularity that linearly drove classification accuracy near its maximal level. Intriguingly, in this period the information held individually by nodes in the first hidden layer about the class increased substantially before ultimately returning to a lower level. This was reflected in changes to low-dimensional embedding distance Within categories (Fig. 4D), which first expanded and subsequently contracted in step with the changes in information held individually by the HL1 nodes about the class. This more complex reconfiguration appeared to organize sets of nodes into modules in a way that involved the most informative nodes about the class becoming segregated hubs within each module and appeared to support the creation of higher order “features” in the next layer. In other words, the reduction in information held by individual nodes in the hidden layer appears to be due to their specialization within modules in carrying this higher-order feature information rather than information directly about the class.

The final period (light purple in Fig. 1C) involved a subsequent reduction in modularity, along with a consolidation that was similarly reflected in an expansion of Between category distance in the low-dimensional embedding of the activity patterns (Fig. 4D). These changes are aligned with a continued increase in information held by the second hidden layer nodes about the class, despite the decrease that occurred earlier in the first hidden layer. Further analysis suggested that this period involved Layer 2 nodes utilizing the higher-order features, combining them (which increases integration/reduces overall modularity) to become more informative about the digit class.

Overall, we conclude that the specialization at earlier layers (i.e., increasing their modularity but decreasing individual information) facilitates the integration of information at later layers, which occurs later in learning. These general trends were confirmed in application to the eMNIST dataset and may represent general learning principles of distributed information processing in networked systems. While some of these features (e.g., increased modularity with learning) are consistent with findings in the systems neuroscience literature, there are others that were more subtle than has been observed in biological systems (e.g., non-topological reconfigurations in early period; late decreases in modularity without performance decrement), and hence may be related to idiosyncrasies inherent within the training of artificial neural networks (e.g., ‘random’ weights at initialisation and back-propagation induced over-fitting, respectively). Future studies will be key to addressing these important open questions.

This study was designed to expose the inner workings of an architecturally basic neural network using a common training dataset. These features were chosen due to their simplicity, which we hoped would help to refine the clarity of the resultant network and informational signatures. With these constraints in mind, a clear open question is whether other more complex network architectures, such as recurrent^44^, convolutional^29^, echo state^45^ or generative adversarial networks^46^, will share similar information processing principles^2,20,21^, or whether the idiosyncratic features that define each of these unique architectures will rely on distinct algorithmic capacities. Since each of these more complex networks use much of the same basic machinery (i.e., diversely connected virtual nodes connected with statistical edges whose weights are trained as a function of trial-and-error), our hypothesis is that most of the informational features described in this study will be pervasive across the family of different network architectures. Regardless, our work provides evidence that network science can provide intuitive explanations for the computational benefits of neural network learning, and helps to highlight the growing intersection between artificial and biological network analyses^47^.

What can network neuroscientists take from the results of this experiment? For one, our observations provide evidence that the tools of systems neuroscience and engineering can indeed be used to understand the function during learning of a complex, high-dimensional system^4,41,42^. Specifically, we observed a substantial increase in network modularity during a simple learning task *in silico*, confirming our *a priori* hypothesis that neural network topology would mirror increases observed in analysis of fMRI networks in human participants^13^. Yet the confirmation can only be considered as partial, as we observed a considerably more subtle temporal partition which, as outlined above, involved three periods that incorporated: edge weight changes without altering the network structure; large increases in modularity to yield seemingly specialist segregated regions; and a regression in overall modularity in later periods. One intriguing possibility is that similar organizing principles underpin learning in biological systems, but are either heavily ingrained in the brain across phylogeny^47^ or occur on sufficiently rapid timescales, such that they are challenging to observe within the limitations inherent within current measurement techniques. Another possibility is that biological brains (and their learning processes) are fundamentally different to what we have observed here, however future experimental work will be required to effectively elucidate these answers.

In addition to these conceptual issues, there are several benefits to analysing neural networks that are not readily apparent in neurobiological analyses. For instance, in the case of feed-forward neural networks, we know the direct, “ground-truth” mapping between inputs and nodes, whereas in neuroscience, this mapping is often opaque. For instance, recordings from standard neuroimaging approaches are often conflated by non-neuronal signals (such as the heartbeat and respiration), and there is also often an indirect mapping between information in the world and the brain’s sensory receptors (e.g., the location of the animal’s gaze can alter the information entering the brain). In addition, standard network approaches in neuroscience require a noisy estimation of network edges, whereas the process here allows us to observe the edge weights directly. It is our hope that these benefits can be used to improve our understanding of the computational benefits of systems-level organizing principles, which in turn can be brought back to neuroscience to accelerate progress in our understanding of the systems-level mechanisms that comprise effective neurological function.

With all that said, neural networks are clearly nowhere near as complex as the typical nervous systems studied by neuroscientists^48^. Most artificial neural networks treat all nodes as identical, whereas diversity reigns supreme in the biological networks of the brain^49^. One key feature of this diversity is the multi-compartment nature of specialist neuronal populations^50^, which render neuronal responses as fundamentally and inextricably non-linear. Crucially, embedding these features into artificial networks has been shown to afford key computational features over more standard approaches^51^. These axonal adaptations are just one of numerous non-linear mechanisms inherent to the brain, including both structural circuit-based mechanisms^52^, as well as more dynamic neuromodulatory gain modulation^53,54^, which are rarely explicitly modelled in neural network studies (though see ^55^). All these features are likely the result of the fact that biological neural networks have been shaped over countless phylogenetic generations, and through this process have inherited innumerable features that have helped to adapt organisms to their different environments^47^. This is in stark contrast to the typical (and admittedly pragmatic) ‘random’ starting point employed in standard neural network experiments – precisely how our psychological and cognitive capacities emerge over the course of phylogeny remains a fascinating source of inspiration for evolving artificial networks^56^. Despite these differences, we maintain that by analysing an artificial network with systems-neuroscience analyses, we can improve our ability to interpret the inner workings of the “black box” of the brain^21^.

In conclusion, we used a systems-level perspective to reveal a series of three serial periods of topological reconfiguration across the course of network training. These periods are associated with distinctive changes to the network and provide important hints as the complex properties that underly efficient distributed information processing in complex systems. Interestingly, many higher-order network properties are shared between the brain and ANNs, suggesting the existence of more general network learning principles that may be uncovered using methods from network science and information theory.

## Methods

### Feed-forward Neural Network

A prototypical, four-layer, feed-forward neural network with no non-linearities was created with randomized weights (edge strengths: −1 to 1). The input layer was designed to take 784 inputs, which themselves were flattened from a 28×28 greyscale pixel array from the MNIST dataset^57^. The input layer was fully connected to a hidden layer of 100 nodes (HL1), which in turn was fully connected to a second hidden layer of 100 nodes (HL2). The second hidden layer was then fully connected to a 10-node output layer. The activation function was a standard sigmoid for the incoming connections at each hidden layer (exponent = 1), and a soft max at the output layer. The maximum value across the nodes of the output layer was taken to reflect the ‘response’ of the network. Each result in our study was also replicated in a separate eMNIST dataset, which was identical to MNIST, but had 26 hand-written letters, as opposed to 10 hand-written digits^58^.

### Training Approach

The network was trained with backpropagation using a Stochastic Gradient Descent optimiser. To aid interpretation, the learnt bias at each neuron was kept to zero and no regularisation was used. The weights and activities were saved as training progressed over the course of a number of epochs (SGD: 100,000). Accuracy was defined as the percentage of trials in a held-out, 10,000 trial testing set in which the maximum value of the output layer was matched with the test category.

### Network Construction

The weighted and signed edges from each asymmetric layer of the neural network were concatenated together to create an asymmetric connectivity matrix. Each connectivity profile was placed in the upper triangle of the matrix (see Fig. 2). To ensure that this step did not adversely affect the topological estimates, each experiment was conducted separately on: a) each layer in turn; b) only the upper triangle of the connectivity matrix. Similar patterns were observed when we re-ran each network separately, suggesting that the embedding did not adversely affect topological interpretation.

### Edge Weight Changes

To determine which edges were associated with maximal change in the first period, we first created a difference score representing the absolute value of edge changes across each pair of epochs in the Early phase. We then calculated the grand mean of this value across the first epoch (i.e., one value for each of the 784 input dimensions, summed across all α edge weights associated with each input node in the Input layer), and then reshaped this vector such that it matched the dimensions of the Input data (i.e., 28^2^ pixels). These values were then compared to the information values (see Mutual Information section below). Correlations between nodes with edge changes across the different periods and IP were compared statistically using an online calculator (https://www.psychometrica.de/correlation.html#dependent)^35^.

### Modularity Maximization

The Louvain modularity algorithm from the Brain Connectivity Toolbox (BCT^73^) was used on the neural network edge weights to estimate community structure. The Louvain algorithm iteratively maximizes the modularity statistic, *Q*, for different community assignments until the maximum possible score of *Q* has been obtained (see Equation 1). The modularity of a given network is therefore a quantification of the extent to which the network may be subdivided into communities with stronger within-module than between-module connections.

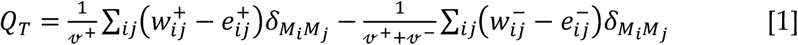

where *v* is the total weight of the network (sum of all negative and positive connections), *w_ij_* is the weighted and signed connection between nodes *i* and *j*, *e_ij_* is the strength of a connection divided by the total weight of the network, and *δ_MiMj_* is set to 1 when nodes are in the same community and 0 otherwise. ‘+’ and ‘–‘ superscripts denote all positive and negative connections, respectively.

For each epoch, we assessed the community assignment for each region 500 times and a consensus partition was identified using a fine-tuning algorithm from the BCT. We calculated all graph theoretical measures on un-thresholded, weighted and signed undirected, asymmetric connectivity matrices^73^. The stability of the γ parameter (which defines the resolution of the community detection algorithm) was estimated by iteratively calculating the modularity across a range of γ values (0.5-2.5; mean Pearson’s r = 0.859 +− 0.01) on the time-averaged connectivity matrix for each subject – across iterations and subjects, a γ value of 1.0 was found to be the least variable, and hence was used for the resultant topological analyses. A consensus clustering partition was defined across all epochs using *consensus_und.m* from the BCT. The resultant solution contained 10 clusters that each contained nodes that were distributed across multiple layers (i.e., Input, HL_1_, HL_2_ and Output).

### Cartographic Profiling

Based on time-resolved community assignments, we estimated within-module connectivity by calculating the time-resolved module-degree Z-score (MZ; within module strength) for each region in our analysis (Equation 2)^74^, where κiT is the strength of the connections of node *i* to other nodes in its module si at time *T*, 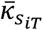 is the average of κ over all the regions in s*i* at time *T*, and 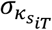 is the standard deviation of κ in si at time *T*.

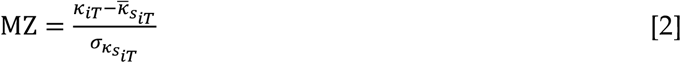

The participation coefficient, *PC*, quantifies the extent to which a node connects across all modules (i.e. between-module strength) and has previously been used to successfully characterize hubs within brain networks (e.g. see ^75^). The PC for each node was calculated within each temporal window using Equation 3, where κisT is the strength of the positive connections of node *i* to nodes in module *s* at time *T*, and κiT is the sum of strengths of all positive connections of nodes *i* at time *T*. Consistent with previous approaches in neuroscience^11,59^, negative connections were removed prior to calculation. The participation coefficient of a region is therefore close to 1 if its connections are uniformly distributed among all the modules and 0 if all of its links are within its own module.

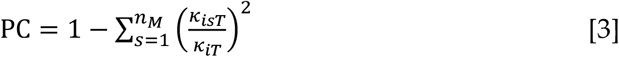

### Mutual Information

We calculated three separate Information measures. To calculate the Information content within each pixel (I*_P_*), we binarized the pixel activity across the 60,000 items from the training set, with a threshold that varied across each pixel so as to maximize the mutual information (MI) that the binarized pixel provides about the class, and then calculated the information within each pixel: MI(pixel,class). To calculate the Information content I*_D_* = MI(pixelOn,class) within each pixel when the pixel was active (after thresholding), we averaged the pointwise MI for each training item, 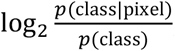, only over the items where the pixel was on (pixelOn). Note that I*_P_* and I*_D_* were negatively correlated across the 28×28 input dimension (r = −0.560, *p_PERM_* < 0.0001), suggesting that the total information from the pixel is dominated by samples when the pixel is inactive (which would be correspondingly averaged as MI(pixelOff,class)). To calculate the Information content within each hidden layer node (I*_H_*), we calculated the mutual information for each node (binarized at activity = 0.5) with the digit class. All MI values were computed using the open source JIDT software^60^.

### Principal Components Analysis

Activity values from the test trials from the input, HL1 and HL2 layers from each epoch were concatenated to form a multi-epoch time series. The data were normalized and then a spatial PCA was performed on the resultant data^41^. The top 3 eigenvectors were used to track the data within a low-dimensional embedding space (Fig. 3), and the percentage explained variance was tracked across all learning epochs. The eigenvectors from the concatenated data were then used to estimate the leading eigenvalues across all training epochs. The analysis was also re-run with activity patterns in HL1 and HL2 separately (i.e., independent of the input layer; Fig. S4). The average value for each exemplar was then used to create two distance measures: Between-category distance, which was defined as the average between-category Euclidean distance at each epoch; and Within-category distance, which was defined as the average within-category Euclidean distance within each epoch.

### Permutation Testing

We used non-parametric testing to determine statistical significance of the relationships identified across our study^61^. A distribution of 10,000 Pearson’s correlations was calculated for each comparison, against which the original correlation was compared. Using this approach, the *p*-value was calculated as the proportion of the null distribution that was less extreme than the original correlation value. In many instances, the effects we observed were more extreme than the null distribution, in which case the *p*-value was designated as *p_PERM_* < 0.0001.

## Acknowledgements

We would like to thank Tim Verstynen, Richard Gao and Manoj Thomas for their feedback on the manuscript.

## Notes

### Competing Interest Statement

The authors have declared no competing interest.

